# Strain-dependent inhibition of *Clostridioides difficile* by commensal *Clostridia* encoding the bile acid inducible *(bai)* operon

**DOI:** 10.1101/2020.01.22.916304

**Authors:** A.D. Reed, M.A. Nethery, A. Stewart, R. Barrangou, C.M. Theriot

## Abstract

*Clostridioides difficile* is one of the leading causes of antibiotic-associated diarrhea. Gut microbiota-derived secondary bile acids and commensal *Clostridia* that encode the bile acid inducible (*bai*) operon are associated with protection from *C. difficile* infection (CDI), although the mechanism is not known. In this study we hypothesized that commensal *Clostridia* are important for providing colonization resistance against *C. difficile* due to their ability to produce secondary bile acids, as well as potentially competing against *C. difficile* for similar nutrients. To test this hypothesis, we examined the ability of four commensal *Clostridia* encoding the *bai* operon (*C. scindens* VPI 12708, *C. scindens* ATCC 35704, *C. hiranonis*, and *C. hylemonae*) to convert CA to DCA *in vitro*, and if the amount of DCA produced was sufficient to inhibit growth of a clinically relevant *C. difficile* strain. We also investigated the competitive relationship between these commensals and *C. difficile* using an *in vitro* co-culture system. We found that inhibition of *C. difficile* growth by commensal *Clostridia* supplemented with CA was strain-dependent, correlated with the production of ∼2 mM DCA, and increased expression of *bai* operon genes. We also found that *C. difficile* was able to outcompete all four commensal *Clostridia* in an *in vitro* co-culture system. These studies are instrumental in understanding the relationship between commensal *Clostridia* and *C. difficile* in the gut, which is vital for designing targeted bacterial therapeutics. Future studies dissecting the regulation of the *bai* operon *in vitro* and *in vivo* and how this affects CDI will be important.

**Importance:** Commensal *Clostridia* encoding the *bai* operon such as *C. scindens* have been associated with protection against CDI, however the mechanism for this protection is unknown. Herein, we show four commensal *Clostridia* that encode the *bai* operon effect *C. difficile* growth in a strain-dependent manner, with and without the addition of cholate. Inhibition of *C. difficile* by commensals correlated with the efficient conversion of cholate to deoxycholate, a secondary bile acid that inhibits *C. difficile* germination, growth, and toxin production. Competition studies also revealed that *C. difficile* was able to outcompete the commensals in an *in vitro* co-culture system. These studies are instrumental in understanding the relationship between commensal *Clostridia* and *C. difficile* in the gut, which is vital for designing targeted bacterial therapeutics.

## Introduction

*Clostridioides difficile* is an anaerobic, spore forming, toxigenic bacterial pathogen (1). *C. difficile* infection (CDI) is a major cause of antibiotic associated diarrhea and a significant health issue, causing 453,000 infections and is associated with 29,000 deaths and 4.8 billion dollars in excess medical costs a year in the U.S. alone (2). While the current first line treatment of vancomycin can resolve CDI, 20-30% of patients who successfully clear the infection experience recurrence (rCDI) within 30 days, and 40-60% of those who experience one episode of rCDI will experience further recurrences (3, 4). Antibiotic use is a significant risk factor for CDI, as antibiotics alter the gut microbiome, causing a loss of colonization resistance against *C. difficile* (5–7). This alteration of the microbiome also affects the gut metabolome, causing a loss in beneficial metabolites, including secondary bile acids generated by the gut microbiota (6, 8). Many of these secondary bile acids are inhibitory to *C. difficile in vitro* and are associated with protection against CDI in mice and humans (9, 10).

Deoxycholate (DCA) is an abundant secondary bile acid in the gut, with concentrations ranging from 0.03-0.7 mM in the non-antibiotic treated gut (11). DCA is synthesized from the primary bile acid cholate (CA) via a multistep pathway that results in 7α-dehydroxylation of CA (12). The enzymes responsible for this synthesis are encoded in the bile acid inducible (*bai*) operon, which is also capable of synthesizing the secondary bile acid lithocholate (LCA) from the primary bile acid chenodeoxycholate (CDCA) (13). A small population of commensal bacteria encoding the *bai* operon are capable of transforming CA to DCA, including *Clostridium scindens, Clostridium hylemonae* and *Clostridium hiranonis* which are members of *Clostridium* cluster XIVa (14–16). Several enzymes within the *bai* operon are capable of completing the steps in this transformation, including the bile acid transporter BaiG, the bile acid 7α-dehydratase BaiE, and the flavoprotein BaiN (13, 17–19). While regulation of the *bai* operon has yet to be completely elucidated, *in vitro* studies show that CA upregulates genes in the *bai* operon and DCA downregulates them in *C. scindens* ATCC 35704, *C. hylemonae*, and *C. hiranonis* (20, 21).

Secondary bile acids are able to inhibit different stages of the *C. difficile* lifecycle. DCA alone is able to inhibit the outgrowth of *C. difficile*, reduce motility and decrease expression of flagellar proteins and toxins *in vitro* (9, 22–24). *In vivo* studies show the presence of *baiCD*, a gene needed for 7α-dehydroxylation, is negatively correlated with CDI in humans, although in another study *C. scindens* was present in the same stool samples as *C. difficile* (25, 26). In addition, *C. scindens* ATCC 35704 is associated with protection against CDI in mouse models, and *C. hiranonis* is negatively correlated with the presence of *C. difficile* in canines, but the exact mechanism of this potential protective effect is still unknown (27–29).

While the production of inhibitory metabolites such as secondary bile acids may be responsible for these potential protective effects, competition for nutrients from other bacteria in the gut, including commensal *Clostridia*, may also play a role. Nutrient competition is another mechanism by which the gut microbiota provides colonization resistance against pathogens. A decrease in specific gut metabolites that *C. difficile* requires for growth (e.g. proline, branched chain amino acids, and carbohydrates) have been associated with CDI in a mouse model and in humans (30–32). In support of this, colonization of a susceptible host by a non-toxigenic strain of *C. difficile* can protect against later colonization by a toxigenic strain (33, 34). This suggests that colonization by a bacterial strain with similar nutritional requirements can have a protective effect on the host. There is evidence that *C. difficile* shares some nutritional requirements with commensal *Clostridia*, including the amino acid tryptophan and the vitamins pantothenate and pyridoxine, which both *C. scindens* ATCC 35704 and *C. difficile* are auxotrophic for (21, 35). *C. difficile* is also auxotrophic for five additional amino acids other than tryptophan, including isoleucine, leucine and proline which are all highly efficient electron donors or acceptors in Stickland fermentation (35, 36). Products of Stickland fermentation are important for growth in *C. difficile* and many other *Clostridia* such as *Clostridium sticklandii* and *Clostridium sporogenes* (35–40)*. C. hiranonis* and *C. hylemonae* both contain genes encoding for enzymes involved in Stickland fermentation that were highly expressed *in vivo. C. scindens* ATCC 35704 has also demonstrated *in vitro* genomic potential for Stickland fermentation (20, 21). Therefore, these commensal *Clostridia* could potentially compete with *C. difficile* for the amino acids it requires for growth and colonization.

In this study, we hypothesized that commensal *Clostridia* are important for providing colonization resistance against *C. difficile* due to their ability to produce secondary bile acids as well as potentially competing against *C. difficile* for similar nutrients. This hypothesis was tested by examining the ability of four commensal *Clostridia* encoding the *bai* operon (*C. scindens* VPI 12708, *C. scindens* ATCC 35704, *C. hiranonis*, and *C. hylemonae*) to convert CA to DCA *in vitro.* The amount of DCA produced was analyzed and the inhibitory effect of the supernatants against a clinically relevant strain of *C. difficile* was tested. We also investigated the competitive relationship between these commensals and *C. difficile* using an *in vitro* co-culture system. We found that inhibition of *C. difficile* growth by commensal *Clostridia* supplemented with CA was strain-dependent, and correlated with the production of ∼2 mM DCA, and increased expression of *bai* operon genes. We also found that *C. difficile* was able to outcompete all four commensal *Clostridia* in an *in vitro* co-culture system. These studies will be instrumental in understanding the relationship between commensal *Clostridia* and the pathogen *C. difficile* in the gut, which is vital for designing targeted bacterial therapeutics. Future studies dissecting the regulation of the *bai* operon *in vitro* and *in vivo* and how this affects CDI will be important.

## Methods

### Genomic analysis of commensal *Clostridia* strains and the *bai* operon

The *bai* operon alignment was constructed by first extracting the positional information for each *bai* gene of interest from Geneious (41), then obtaining amino acid identity percentage through BLASTp alignments (42) against coding sequences from the reference strain *C. scindens* ATCC 35704. This data was visualized using the publicly available gggenes R package (43) with slight modifications. All vs *C. scindens* ATCC 35704 alignments were visualized using the BLAST Ring Image Generator (44), including entries for GC Content and GC Skew. BLASTn was used for this alignment, with an upper identity threshold of 90%, a lower identity threshold of 50%, and a ring size of 30.

### Bacterial strain collection and growth conditions

The *C. difficile* strain used in this study was R20291, a clinically relevant strain from the 027 epidemic ribotype (9). All assays using *C. difficile* were started from spore stocks, which were prepared and tested for purity as described previously (9, 45). *C. difficile* spores were maintained on brain heart infusion (BHI) media supplemented with 100 mg/L L-cysteine and 0.1% taurocholate (T4009, Sigma-Aldrich). Then cultures were started by inoculating a single colony from the plate into BHI liquid media supplemented with 100 mg/L L-cysteine.

Four strains of commensal *Clostridia* encoding the *bai* operon were used in this study. *Clostridium scindens* ATCC 35704 (Cat #35704) was purchased from the American Type Culture Collection. *Clostridium scindens* VPI 12708, *Clostridium hylemonae* TN-271 and *Clostridium hiranonis* TO-931 were obtained from Jason M. Ridlon (University of Illinois Urbana-Champaign, United States). All strains were maintained on 15% glycerol stocks stored in −80 °C until use and were grown in BHI supplemented with 100 mg/mL L-cysteine. Media for *C. hiranonis* was BHI supplemented with 100 mg/mL L-cysteine and 2 µM hemin. All strains used in this study were grown under 2.5% hydrogen in anaerobic conditions (Coy, USA) at 37 °C.

### Minimum inhibitory concentration (MIC) assay with the addition of bile acids

MICs were determined using the modified Clinical and Laboratory Standards Institute broth microdilution method as described previously (45). The inoculum was prepared by the direct colony suspension method. All cell concentrations were adjusted to an optical density at 600 nm (OD_600_) of 0.01. Briefly, the MIC plates were prepared by making fresh bile acid dilution stocks in the test media, then adding 90 µL to each well such that the final concentration of the bile acid after the addition of cells (10 µL) ranged from 0.04 mM to 10 mM for CA (102897, MP Biomedicals), and 0.01 mM to 2.5 mM for DCA (D6750, Sigma Aldrich). Four biological replicates were performed. Positive controls were inoculated cells with no bile acid in test media. Uninoculated media was used as a control for sterility. *C. scindens, C. hiranonis* and *C hylemonae* were incubated for 48 hr while *C. difficile* was incubated for 24 hr anaerobically at 37 °C. All assays were performed in BHI supplemented with 100 mg/L of L-cysteine. For assays involving *C. hiranonis*, the BHI was also supplemented with 2 µM hemin.

### Growth kinetics assay

Cultures of *C. scindens, C. hiranonis, C. hylemonae* and *C. difficile* liquid cultures were started from a single colony and grown for 14 hr, then sub-cultured 1:10 and 1:5 in liquid media and allowed to grow for 3 hr or until doubling. Cultures were then diluted in fresh BHI supplemented with varying concentrations of CA or DCA so that the starting OD_600_ was 0.01. The growth media for *C. hylemonae* was BHI supplemented with 2 µM hemin and varying concentrations of CA or DCA. Growth studies were performed in 200 µL of media over 24 hr using a Tecan plate reader inside the anaerobic chamber. OD_600_ was measured every 30 min for 24 hr, and the plate was shaken for 90 sec before each reading was taken. Three technical replicates were performed for each concentration of bile acid, and three biological replicates were performed for each organism.

### *C. difficile* inhibition assay with supernatants from commensal *Clostridia* supplemented with and without bile acids

Cultures of *C. scindens* and *C. hiranonis* were grown in fresh BHI supplemented with 0 mM, 0.25 mM or 2.5 mM of CA, while cultures of *C hylemonae* were grown in fresh BHI supplemented with 0 mM, 2.5 mM or 7 mM of CA. After 14 hr of growth, cultures were spun down anaerobically at 6,000 rpm for 5 min. Supernatants were then sterilized under anaerobic conditions using a 0.22 µM filter and used in the inhibition assay at a ratio of 4 parts supernatant to one part BHI.

Cultures of *C. difficile* were started from a single colony and grown for 14 hr, then subcultured 1:10 and 1:5 in liquid media and allowed to grow for 3 hr or until doubling, then diluted to 0.01 OD in a mixture of 4 parts filtered supernatant to 1 part BHI. *C. difficile* grown in a mixture of 4 parts PBS to 1 part BHI was used as a control for this assay.

Bile acid controls included *C. difficile* grown in BHI supplemented with 0.25 mM, 2.5 mM 7 mM CA as well as BHI supplemented with 0.25 mM or 2.5 mM DCA. *C. difficile* grown in BHI without bile acid supplementation was used as a positive control. Cultures were allowed to incubate for 24 hr anaerobically at 37 °C, then dilutions were plated on BHI plates, and the number of colony forming units (CFUs) was calculated the next day. Aliquots of the supernatants from *C. scindens, C. hiranonis* and *C. hylemonae* from this assay were stored at −80 °C for later bile acid metabolomic analysis.

#### Bile acid metabolomic analysis

Culture media supernatants were diluted 1:100 in methanol (2 µL supernatant, 198 µL MeOH) and analyzed by UPLC-MS/MS. The analysis was performed using a Thermo Vanquish LC instrument (Thermo Fisher Scientific, San Jose, CA) coupled to a Thermo TSQ Altis triple quadrupole mass spectrometer (Thermo Fisher Scientific, San Jose, CA) with a heated electrospray ionization (HESI) source. Chromatographic separation was achieved on a Restek Raptor C18 column (2.1 x 50 mm, 1.8 mM) maintained at 50°C. The following linear gradient of mobile phase A (5 mM ammonium acetate) and mobile phase B (1:1 MeOH/MeCN) was used: 0-2 min (35-40%B, 0.5 mL/min), 2-2.5min (40-45%B, 0.5 mL/min), 2.5-3.5 min (45-50%B, 0.5 mL/min), 3.5-4.6 min (50-55%B, 0.5 mL/min), 4.6-5.7 min (55-80%B, 0.8 mL/min), 5.7-5.9 min (80-85%B, 0.8 mL/min), 5.9-6.5 min (85%B, 0.8 mL/min), 6.5-8.5 min (35%B, 0.5 mL/min). For quantification, certified reference material (50 mg/mL) for CA and DCA were obtained from Cerilliant. These individual stocks were combined and diluted to achieve a 250 mM working standard solution. Seven calibration standards ranging from 8 nM to 125 mM were prepared by serially diluting the working standard solution. Both samples and standards were analyzed (2 mL injections) in negative ion mode (spray voltage 2.5 kV, ion transfer tube temperature 325°C, vaporizer temperature 350°C, sheath gas 50 a.u., aux gas 10 a.u., sweep gas 1 a.u.) using a Q1 resolution of 0.7 m/z, a Q3 resolution of 1.2 m/z and a collision energy of 22 V. The following multiple reaction monitoring (MRM) transitions were used: 407.1 → 407.1 (CA), 391.1 → 391.1 (DCA).

#### Targeted data processing

Peak integration and quantification were performed in TraceFinder 4.1 (Thermo Fisher Scientific, San Jose, CA). Individual standard curves for CA and DCA were constructed using peak areas from the quantifier transitions (407.1 → 407.1 for CA, 391.1 → 391.1 for DCA). The concentrations of CA and DCA in the study samples were calculated in an identical manner relative to the regression line. Calibration curves for CA and DCA had R^2^ values ranging of 0.9995 and 0.9983, respectively, for the linear range of 8 nM to 125 mM with a weighting of 1/x. Validation of the curve with a QC sample (2.5 mM) passed with a threshold of 10%.

### RNA extraction from commensal *Clostridia* cultures supplemented with CA

*C. scindens, C. hiranonis* and *C. hylemonae* liquid cultures were started from a single colony and grown for 14 hr, then subcultured 1:10 and 1:5 in liquid media and allowed to grow for 3 hr or until doubling. Cultures of *C. scindens* and *C. hiranonis* were then diluted to 0.1 OD in fresh BHI supplemented with 0 mM, 0.25 mM or 2.5 mM of CA, while cultures of *C hylemonae* were diluted to 0.1 OD in fresh BHI supplemented with 0 mM, 2.5 mM or 7 mM of CA. Cultures were allowed to grow to mid log phase (OD of 0.3-0.5) then half of the culture was removed and stored for later extraction. The remaining portion of the culture was allowed to grow for 14 hr until stationary phase was reached.

Cultures were fixed by adding equal volumes of a 1:1 mixture of EtOH and acetone and stored in the −80 °C for later RNA extraction. For extraction, the culture was thawed then centrifuged at 10,000 rpm for 10 min at 4 °C. The supernatant was discarded and the cell pellet resuspended in 1 mL of 1:100 BME:H_2_O, then spun down at 14,000 rpm for 1 min. The cell pellet was resuspended in 0.3 mL of lysis buffer from the Ambion RNA purification kit (AM1912, Invitrogen) then sonicated while on ice for 10 pulses of 2 sec with a pause of 3 sec between each pulse. Extraction was then performed following the manufacturers protocol from the Ambion RNA purification kit.

### Reverse transcription and quantitative real-time PCR

Reverse transcription and quantitative real-time PCR was performed as described previously (30). Briefly, RNA was depleted by using Turbo DNase according to the manufacturer’s instructions (AM2238, Invitrogen. The DNased RNA was then cleaned using an RNA clean up kit (R1019, Zymo) according to manufacturer instruction and DNA depletion was verified by amplifying 1 µL of RNA in a PCR reaction. The DNA depleted RNA was used as the template for reverse transcription performed with Moloney murine leukemia virus (MMLV) reverse transcriptase (M0253, NEB). The cDNA samples were then diluted 1:4 in water and used in quantitative real-time PCR with gene-specific primers using SsoAdvanced Universal Sybr green Supermix (1725271, Bio-Rad) according to the manufacturer’s protocol. Amplifications were performed in technical quadruplicate, and copy numbers were calculated by the use of a standard curve and normalized to that of a housekeeping gene. *gyrA* was the housekeeping gene used for *C. scindens* and *rpoC* was the housekeeping gene used for *C. hiranonis* and *C. hylemonae*.

The housekeeping gene for each strain was determined by testing a list of genes (Table 1 and Table 2) using cDNA standardized to a concentration of 0.3 ug/µL. Three technical replicates were performed for each assay, and three biological replicates were performed. *C. scindens* and *C. hiranonis* were tested with RNA from cultures grown to mid log and stationary phase in media supplemented with 0 mM, 0.25 mM or 2.5 mM of CA, while *C. hylemonae* was tested with RNA from cultures grown to mid log and stationary in media supplemented with 0 mM, 2.5 mM or 7 mM of CA, and copy numbers were calculated by the use of a standard curve. Analysis of copy numbers was performed with Normfinder (46) and the gene with the lowest inter-group variance, or stability value was selected (Table 3). The Log_2_ fold change for each gene and each condition (Table S1) was calculated by dividing the expression when CA was added by the expression of the negative control (0 mM CA).

**Table 1:** Candidate Reference Genes

| Putative Housekeeping Gene | Function |
| --- | --- |
| *rpoC* | RNA polymerase subunit C |
| *gyrA* | Gyrase subunit A |
| *gluD* | Glutamate dehydrogenase |
| *adk* | Adenylate kinase |
| *dnaG* | DNA primase |
| *recA* | Recombination protein A |
| *rpsJ* | 30S ribosomal protein S10 |

**Table 2:**
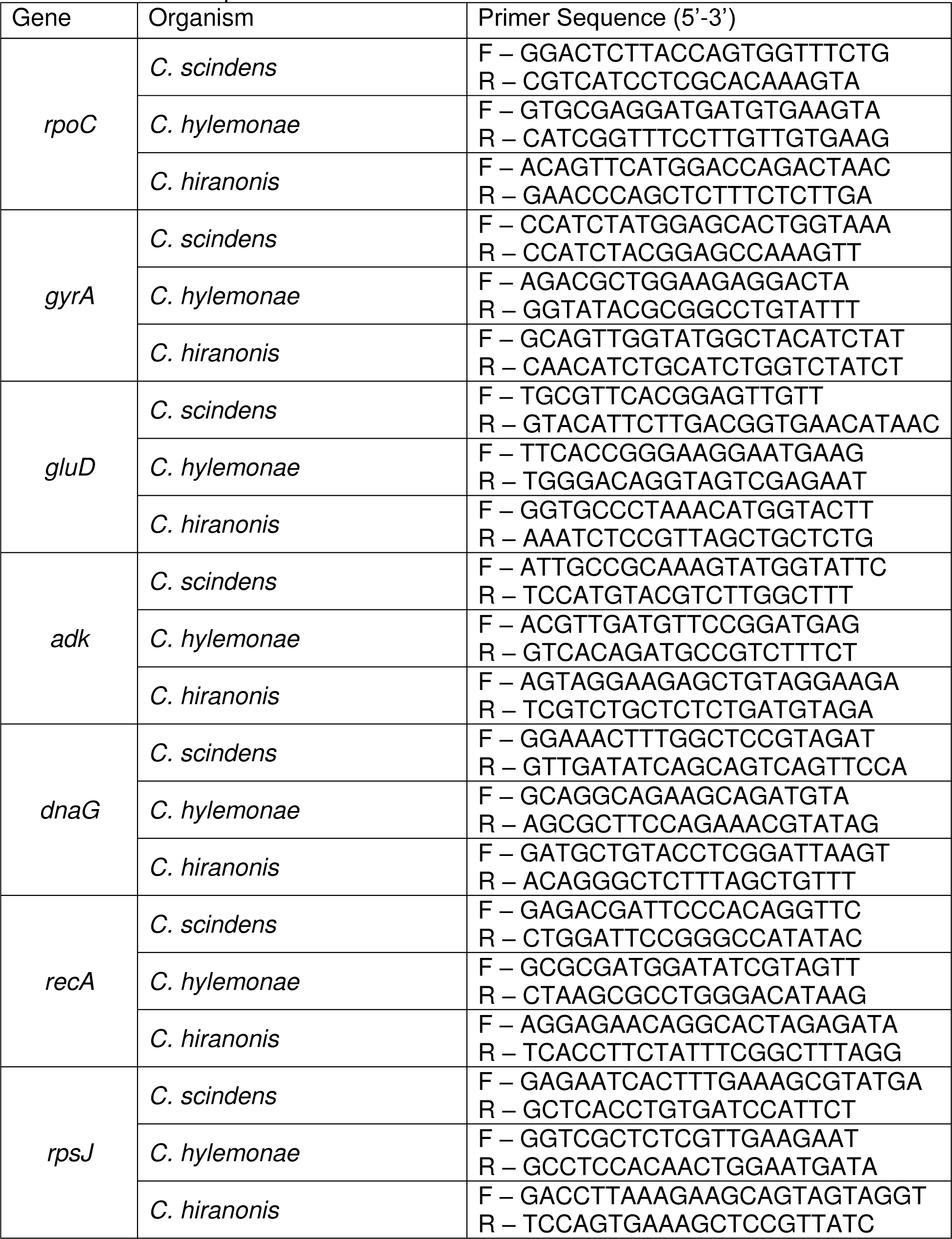
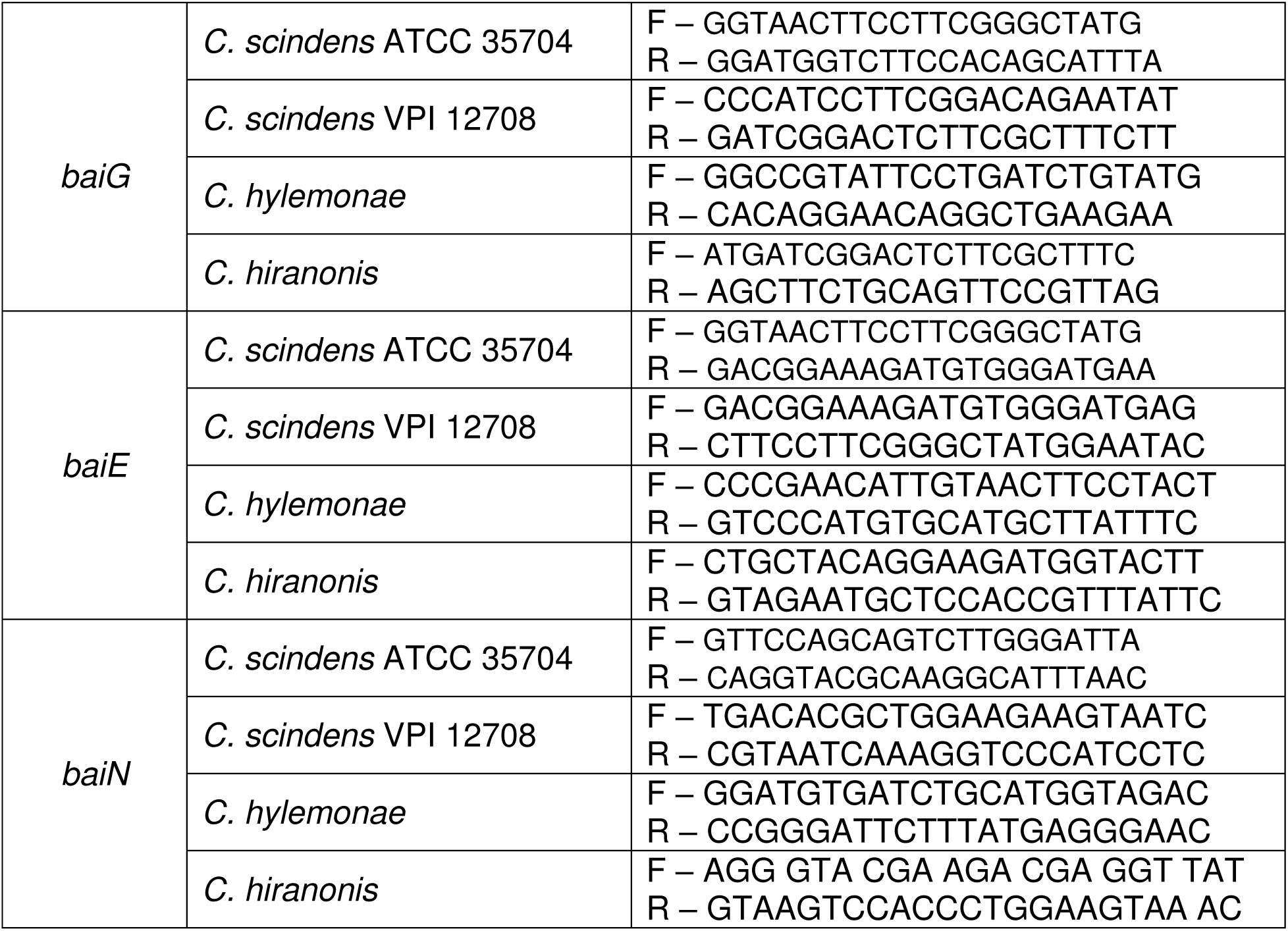
Primer Sequences Used

**Table 3:** Normfinder Analysis

| *C. scindens* ATCC 35704 |  | *C. scindens* VPI 12708 |  | *C. hylemonae* |  | *C. hiranonis* |  |
| --- | --- | --- | --- | --- | --- | --- | --- |
| Gene | Stability Value | Gene | Stability Value | Gene | Stability Value | Gene | Stability Value |
| ***gyrA*** | 0.41 | ***gyrA*** | 0.70 | ***rpoC*** | 0.63 | ***rpoC*** | 0.34 |
| *recA* | 0.45 | *recA* | 0.73 | *gyrA* | 0.65 | *dnaG* | 0.38 |
| *rpoC* | 0.50 | *dnaG* | 0.84 | *dnaG* | 0.89 | *gyrA* | 0.44 |
| *dnaG* | 0.59 | *rpoC* | 1.10 | *recA* | 1.16 | *Adk* | 0.47 |
| *adk* | 0.94 | *adk* | 1.40 | *gluD* | 1.21 | *gluD* | 0.48 |
| *rpsJ* | 0.99 | *rpsJ* | 1.55 | *adk* | 1.54 | *recA* | 0.52 |
| *gluD* | 2.04 | *gluD* | 2.09 | *rpsJ* | 1.79 | *rpsJ* | 0.56 |

### Competition studies between *C. difficile* and commensal *Clostridia*

*C. scindens, C. hylemonae* and *C difficile* liquid cultures were started from a single colony as described above. Monocultures of each organism were diluted to ∼1×10^5^ CFU/mL in fresh BHI media, and 1:1 competition assays were ∼1×10^5^ CFU/mL of each organism in fresh media. Determination of the conversion from OD to CFU/mL was performed for *C. difficile, C. scindens* and *C. hylemonae* using a growth curve to measure OD and CFU/mL. The conversion from OD to CFU/mL was calculated using a linear regression analysis (Figure S1).

Dilutions were performed at 0 hr and after 24 hr of incubation, and colonies were counted to determine the number of CFUs per mL. Colonies were counted after the plates had been incubated long enough to allow individual colonies to form, which was 24 hr for *C. difficile* and 48 hr for *C. scindens* and *C. hylemonae*. Differences in colony morphology and growth time were used to distinguish between *C. scindens*, *C. hylemonae* and *C. difficile*.

### Statistical analyses

Statistical tests were performed using Prism version 7.0c for Mac OS X (GraphPad Software, La Jolla, CA, United States). Significance was determined by using one-way ANOVA to determine significance across all conditions, with Tukey’s test used to correct for multiple comparisons. Statistical significance was set at a p value of < 0.05 for all analyses, (*p < 0.05, **p < 0.01, *** p < 0.001, **** p < 0.0001). All assays were performed with at least three biological replicates.

## Results

### Genomic comparison between commensal *Clostridia* that encode the *bai* operon

The *bai* operon of each strain was extracted for visual alignment and amino acid comparison against reference strain *C. scindens* ATCC 35704 (Figure 1). The operons of both *C. scindens* ATCC 35704 and *C. scindens* VPI 12708 are architecturally similar with each gene within the operon sharing at least 97% identity at the amino acid level. Interestingly, *C. hylemonae* TN 271 possesses a shorter *bai* operon due to the lack of the *baiA2* gene, while the *bai* operon of *C. hiranonis* TO 931 is ∼ 1 kb longer than the reference operon due to expanded intergenic regions. *C. hylemonae* TN 271 shares between 63% and 89% identity with the reference operon and *C. hiranonis* displays between 48% and 90% identity. Notably, the outermost protein coding sequences in the *bai* operon of *C. hylemonae* TN 271 and *C. hiranonis* TO 931, *baiB* and *baiI*, exhibit a significantly reduced percent identity when compared to other genes in the operon.

**Figure 1:**
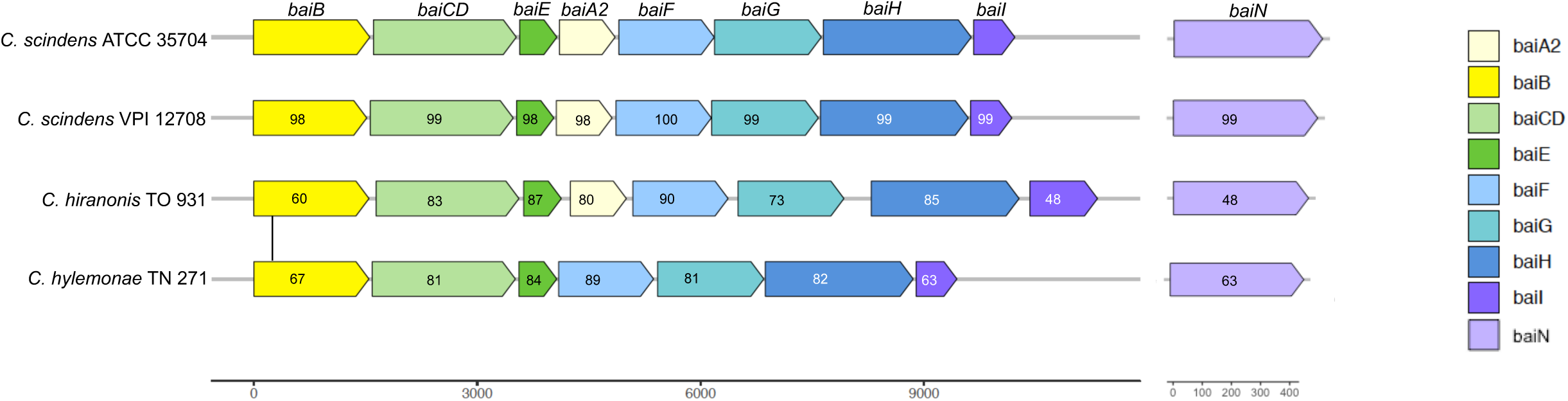
Genomic variation in the *bai* operon encoded by commensal *Clostridia*. *bai* operon alignment across *Clostridium* strains. Each protein sequence was compared against its counterpart in the reference strain *Clostridium scindens* ATCC 35704, generating the amino acid percent identity labeled within each gene.

The differences in identity across these four operons is largely representative of the whole-genome nucleotide comparison of each strain (Figure S2), with *C. hiranonis* TO931 sharing the least identity to the reference strain *C. scindens* ATCC 35704, followed by the slightly higher percent nucleotide identity of *C. hylemonae* TN271, with *C. scindens* VPI 12708 sharing the most nucleotide identity.

### *C. difficile* is more resistant to cholate and less resistant to deoxycholate compared to commensal *Clostridia*

Since secondary bile acids such as DCA are made by specific commensal *Clostridia* encoding the *bai* operon from CA and are inhibitory against *C. difficile*, we sought to examine the resistance profiles of these bacteria with CA and DCA (9). We did this by performing minimum inhibitory concentration (MIC) assays with CA and DCA for *C. difficle* R20291 as well as four commensal *Clostridia* strains that encode the *bai* operon (14–16). *C. difficile* had a high MIC of 10 mM with CA, (Figure 2, Table S1), but a much lower MIC of 1.56 mM with DCA. Of the commensal *Clostridia, C. hylemonae* had the highest MIC against CA with 7 mM, while *C. hiranonis* and the two *C. scindens* strains all had a MIC of 2.5 mM for CA. Of the commensals, *C. hiranonis* was most sensitive to DCA, with a MIC of 0.78 mM. *C. hylemonae* was also sensitive to DCA, with a MIC of 1.25 mM. The two *C. scindens* strains were more resistant to DCA, with *C. scindens* VPI 12708 having a MIC of 1.88 mM and *C. scindens* ATCC 35704 having a MIC of 2.19 mM. Although the commensal *Clostridia* tested have all shown the ability to produce DCA *in vitro*, they all display sensitivity to it. Out of the four commensal strains, only the two *C. scindens* strains had higher resistance to DCA than *C. difficile* R20291. Different concentrations of CA and DCA were also able to alter the growth kinetics of the commensal *Clostridia* and *C. difficile* as seen in Figures S3 and S4.

**Figure 2:**
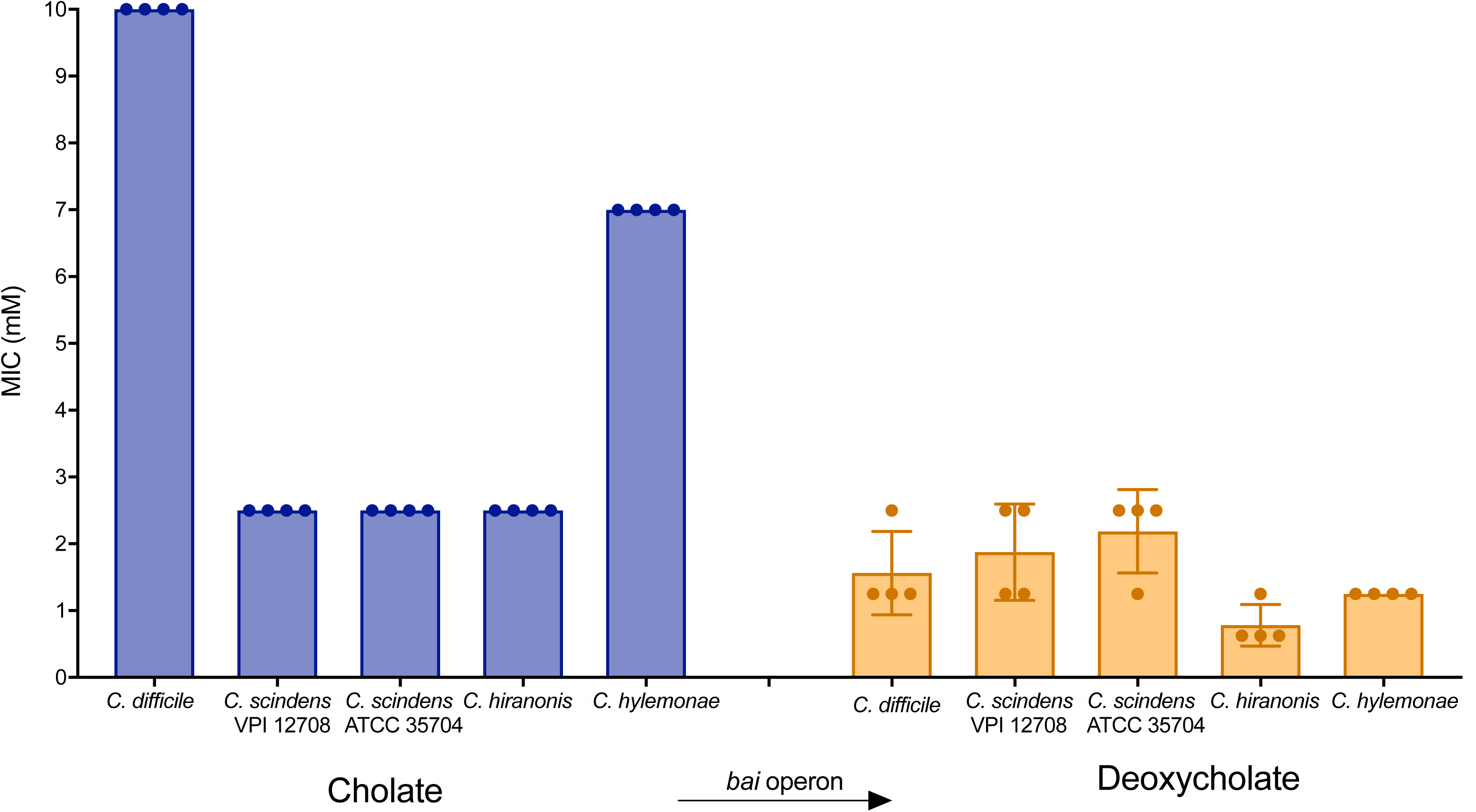
*C. difficile* and *C. hylemonae* are more resistant to cholate than other commensal *Clostridia* tested. The Minimum Inhibitory Concentration (MIC) of CA and DCA was tested on *C. difficile*, *C. scindens* VPI 12708, *C scindens* ATCC 35704, *C. hiranonis* and *C. hylemonae.* The MIC was defined as the lowest concentration of compound that showed no visible growth. Optical density was defined at 24 hr for *C. difficile* and 48 hr for commensal *Clostridia.* Four biological replicates were performed.

### Commensal *Clostridia* inhibit *C. difficile* in a strain-dependent manner that correlates with the conversion of cholate to deoxycholate

While all four commensal strains examined in this study have been shown to produce DCA from CA *in vitro*, we wanted to ascertain whether this was sufficient to inhibit *C. difficile* growth (15, 16, 47). We developed an *in vitro* inhibition assay using supernatants from overnight cultures of commensal *Clostridia* supplemented with and without different concentrations of CA. The supernatants were then added to fresh *C. difficile* cultures to investigate if commensal *Clostridia* capable of producing DCA from CA were able to inhibit *C. difficile* growth. When the supernatant from *C. scindens* VPI 12708 cultures supplemented with 0.25 mM of CA and no CA were added to *C. difficile* cultures, there was no inhibition of growth after 24 hr. When *C. scindens* VPI 12708 cultures were supplemented with 2.5 mM of CA, the supernatant significantly inhibited *C. difficile* growth (Figure 3A). The level of inhibition was similar to the level of inhibition seen when *C. difficile* was grown with 2.5 mM of DCA alone (Figure S5). In order to determine the levels of CA and DCA present in the supernatants added to *C. difficile* cultures, targeted liquid chromatography-mass spectrometry (LC/MS) was performed. *C. scindens* VPI 12708 was able to convert almost all of the CA present in the media to DCA (Figure 3A). When the media was supplemented with 0.25 mM of CA, 0.24 ± 0.02 mM DCA was produced, and 1.97 ± 0.21 mM DCA was produced when the media was supplemented with 2.5 mM CA. Supernatant from *C. scindens* ATCC 35704 cultures also greatly inhibited *C. difficile* when grown in media supplemented with 2.5 mM of CA (Figure 3B), but some inhibition was also seen with supernatant from cultures supplemented with 0.25 mM CA and no CA. This is likely due to the production of tryptophan-derived antimicrobials produced by this *C. scindens* strain that have previously been shown to inhibit *C. difficile* (48). *C. scindens* ATCC 35704 also converted most of the CA present in the media to DCA (Figure 3B) with 0.19 ± 0.05 mM of DCA being produced when the media was supplemented with 0.25 mM CA, and 1.95 ± 0.38 mM DCA being produced when supplemented with 2.5 mM CA. *C. hiranonis* and *C. hylemonae* culture supernatants did not significantly inhibit *C. difficile* growth (Figure 3C and D), regardless of the amount of CA present in the culture media. *C. hiranonis* did convert some of the CA to DCA (Figure 3C), with 0.14 ± 0.03 mM DCA being produced when supplemented with 0.25 mM CA, and 0.78 ± 0.05 mM DCA produced when supplemented with 2.5 mM CA. *C. hylemonae* did not convert any of the CA present in the media to DCA (Figure 3D).

**Figure 3:**
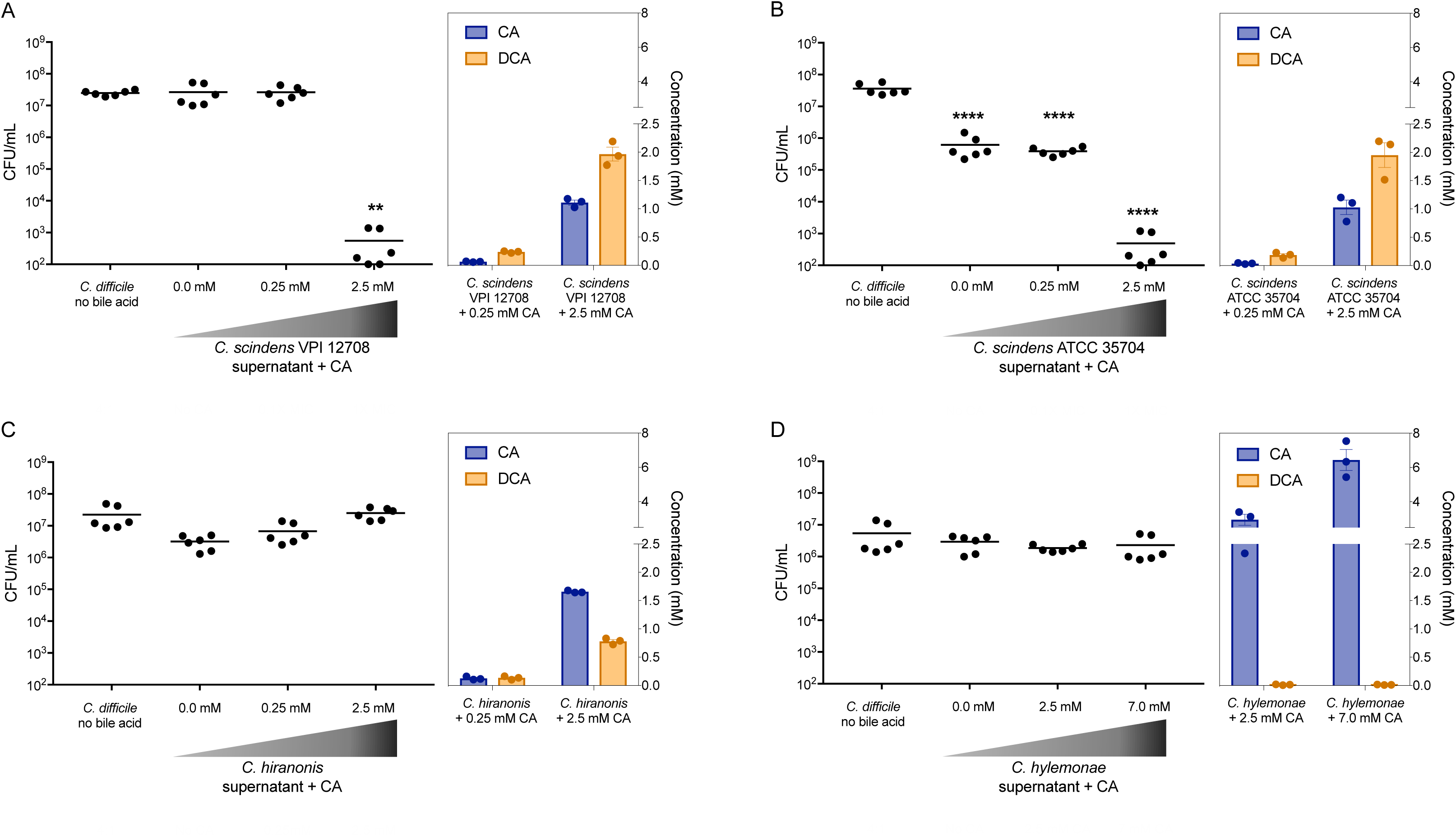
Inhibition of *C. difficile* by *C. scindens* grown in media supplemented with 2.5 mM cholate is correlated with high levels of deoxycholate production. Inhibition of *C. difficile* after 24 hr of growth with supernatants from (A) *C. scindens* VPI 12708, (B) *C. scindens* ATCC 35704, and (C) *C. hiranonis* grown without bile acid supplementation or supplemented with 0.25 or 2.5 mM of CA, and (D) *C. hylemonae* grown without bile acid supplementation or supplemented with 2.5 mM or 7.0 mM of CA. The concentration of CA and DCA in each supernatant is shown to the right of the inhibition data. Experiments were run in duplicate, and three biological replicates were performed. Inhibition by the supernatants was compared to a no bile acid *C. difficile* control consisting of a 4:1 dilution of PBS to BHI. Statistical significance between treatments and the control was determined using one-way ANOVA with Tukey used for multiple comparisons (*, p<0.05; **, p<0.01; ***, p<0.001; ****, p<0.0001).

### Expression of *baiE* and *baiG* is increased when cholate is converted to deoxycholate

To investigate the effect of different concentrations of CA on expression of the *bai* operon, three genes were selected for qRT-PCR analysis. We selected *baiG*, which is responsible for transport of CA into the cell, *baiE*, which is responsible for the irreversible and rate limiting step, and *baiN*, which is capable of performing the first two reductive steps in the pathway (18, 19, 49). Commensal *Clostridia* cultures were grown in a rich media supplemented with different concentrations of CA for a 24 hr period, followed by RNA extraction before qRT-PCR analysis. *C. scindens* VPI 12708 had a significant increase in expression of *baiE* and *baiG* when 0.25 mM or 2.5 mM of CA was present in cultures (Figure 4A). *C. scindens* ATCC 35704 also showed a significant increase in expression in *baiE* and *baiG* when 2.5 mM of CA was present in cultures, but not with 0.25 mM of CA (Figure 4B). *C. hiranonis* had a significant increase in expression in *baiE* and *baiG* when 2.5 mM of CA was present in cultures (Figure 4C), but the increased expression is approximately ten fold less than either *C. scindens* strain with an equal amount of CA in the media. *C. hylemonae* had a decrease in expression of *baiE* when 2.5 mM CA was added (Figure 4D), and decreased expression of *baiE* and *baiG* when 7 mM of CA was added. Expression of *baiN* was not affected by the different CA concentrations in the media, although a small but significant increase was seen in *C. scindens* ATCC 37704 when supplemented with 2.5 mM CA.

**Figure 4:**
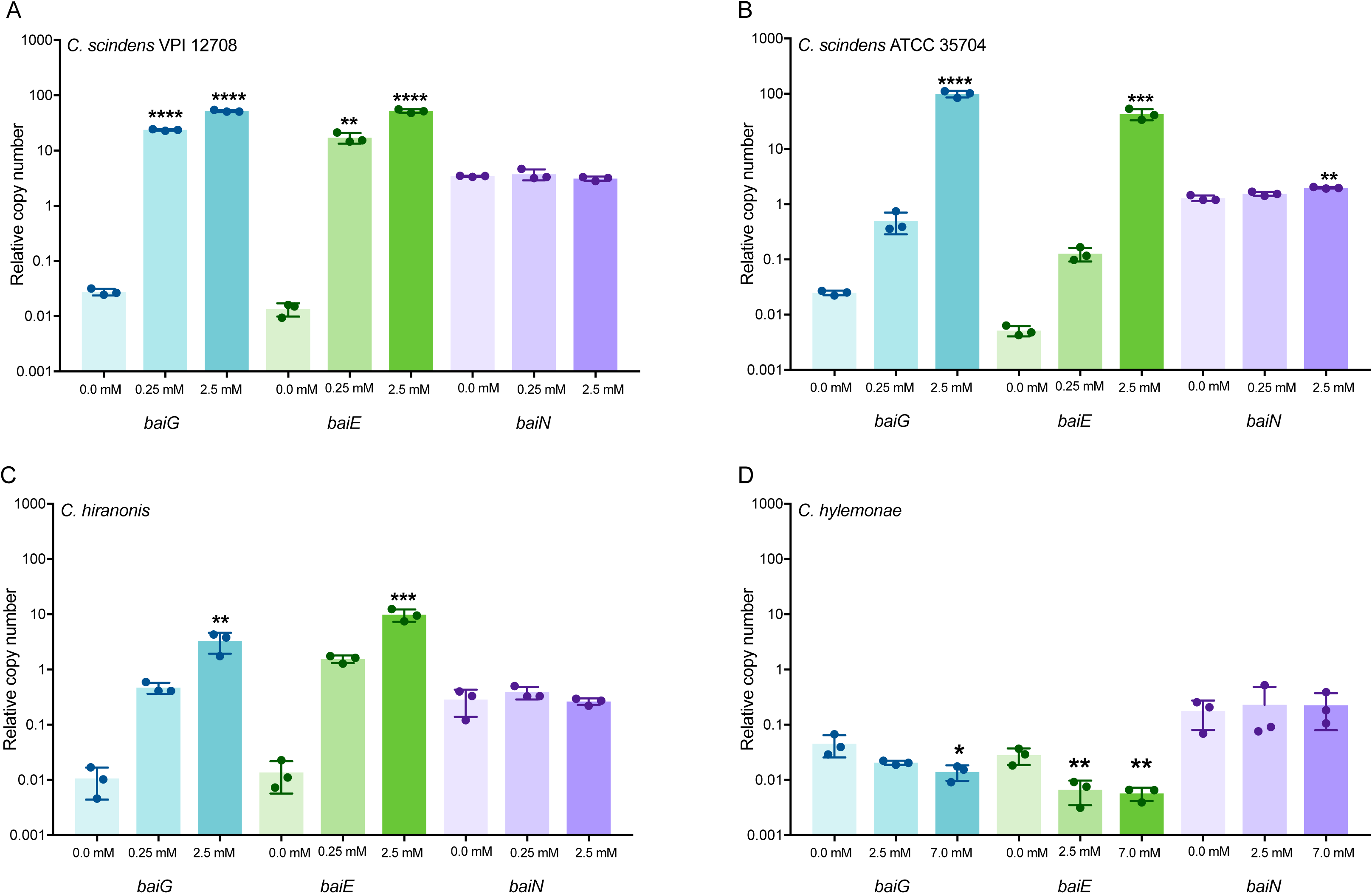
*C. scindens* and *C. hiranonis* have increased expression in *bai* operon genes when media is supplemented with 2.5 mM CA. Expression of *baiG, baiE* and *baiN* in (A) *C. scindens* ATCC 35704, (B) *C. scindens* VPI 12708, and (C) *C. hiranonis* in media without CA or media supplemented with 0.25 mM or 2.5 mM CA, and (D) *C. hylemonae* in media without CA or media supplemented with 2.5 mM or 7.0 mM CA. Experiments were run in quadruplicate, and three biological replicates were performed. Expression in media supplemented with CA was compared to expression in media without CA. Statistical significance was determined by one-way ANOVA with Tukey used for multiple comparisons (*, p<0.05; **, p<0.01; ***, p<0.001; ****, p<0.0001).

### *C. difficile* outcompetes commensal *Clostridia* in a strain dependent manner

In order to explore our additional hypothesis that commensal *Clostridia* are able to compete against *C. difficile* for similar nutrients, we performed 1:1 competition assays between commensal *Clostridia* and *C. difficile* in rich media without the addition of CA (Figure 5 and Figure S6). A monoculture of each strain was used as a control for each replicate performed. The competition index (Figure 5) was calculated by dividing the CFU of each strain after 24 hr of growth in the 1:1 competition assay with the CFU of the monoculture control after 24 hr. The raw data from the competition assays is available in Figure S6. All commensal *Clostridia* growth was significantly inhibited in co-culture with *C. difficile*, with *C. hiranonis* being affected the most. *C. difficile* growth was not negatively affected by any commensal strain except for *C. scindens* ATCC 35704, which is consistent with the inhibition observed by *C. scindens* ATCC 35704 culture supernatant without CA in Figure 3A and the inhibition observed by Kang et al. (48).

**Figure 5:**
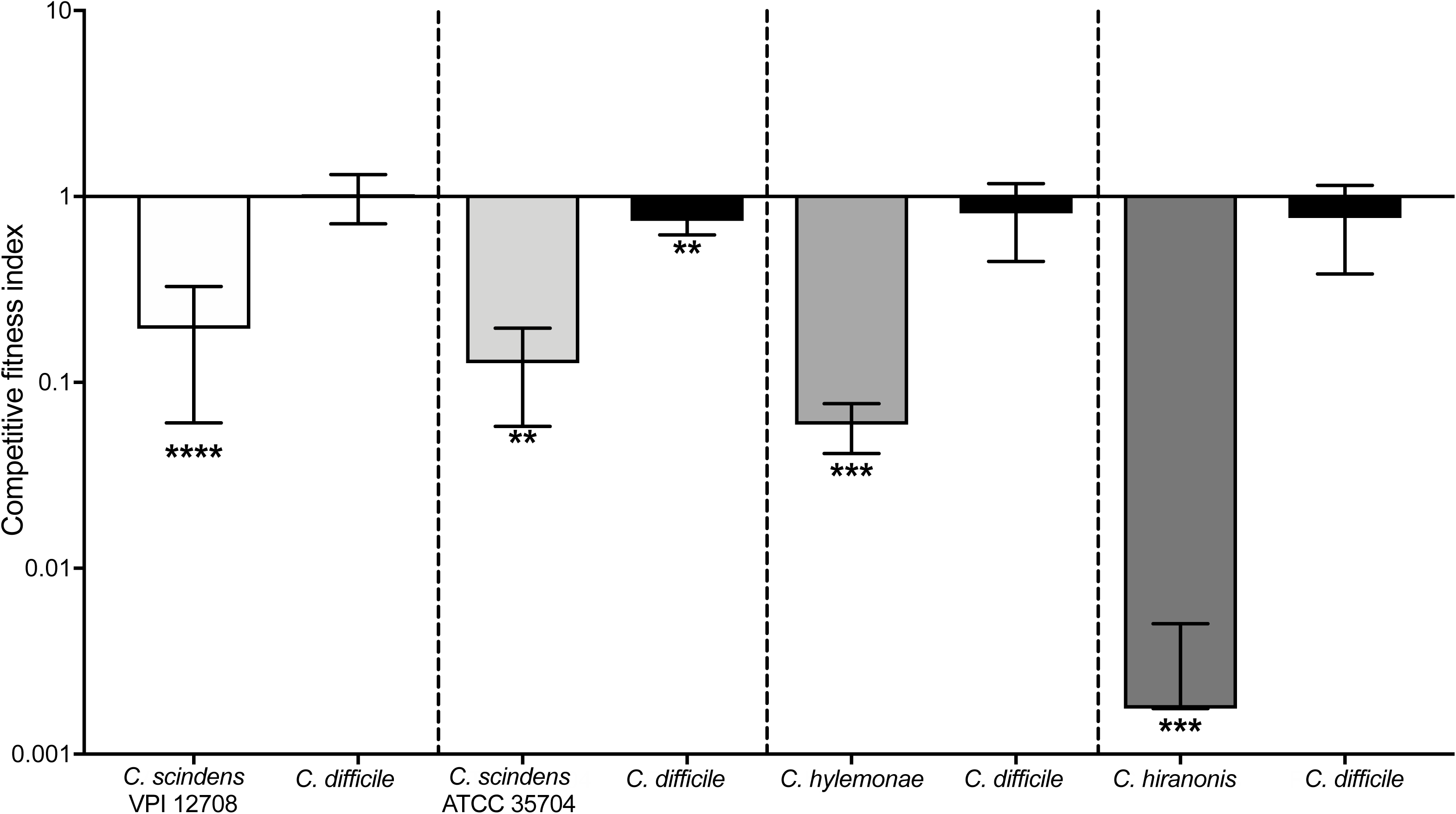
*C. difficile* outcompetes commensal *Clostridia in vitro*. Competition index for 1:1 competition between *C. difficile* and *C. scindens* ATCC 35704, *C. scindens* VPI 12708, *C. hylemonae* and *C. hiranonis.* The competition index value was determined by comparing the CFU/mL of the competition co-culture to the monoculture for each strain at 24 hr. Statistical significance was determined by using students T test (*, p<0.05; **, p<0.01; ***, p<0.001; ****, p<0.0001).

## Discussion

In this study we examined the genomic variation between four commensal *Clostridia* strains containing the *bai* operon, determined their ability to produce DCA when supplemented with CA, and investigated their ability to inhibit *C. difficile in vitro*. While *C. scindens* VPI 12708, *C. scindens* ATCC 35704 and *C. hiranonis* all produced DCA under these conditions (Figure 4), only *C. scindens* strains were able to inhibit *C. difficile* when supplemented with 2.5 mM CA. This is likely due to the efficient conversion of CA to DCA produced when they were supplemented with 2.5 mM CA. *C. hiranonis* produced less DCA (0.78 mM) when supplemented with 2.5 mM CA. This could be due to the lower MIC of *C. hiranonis* with DCA (0.78 mM) compared to *C. scindens* VPI 12708 (1.88 mM) and *C. scindens* ATCC 35704 (2.19 mM). Additionally, *C. difficile* inhibited the growth of all four commensals tested in a 1:1 *in vitro* competition assay without the presence of CA, with only *C. scindens* ATCC 35704 affecting *C. difficile* growth, which is likely due to the antimicrobial activity it produces (48).

Genomically the *bai* operon of *C. scindens* VPI 12708 had a high amino acid similarity (∼97%) to *C. scindens* ATCC 35704, but *C. hylemonae* and *C. hiranonis* were more divergent than expected. This pattern continued when examining the entire genome, with *C. hiranonis* and *C. hylemonae* diverging from *C. scindens* ATCC 35704 more than *C. scindens* VPI 12708. Of particular interest is the lack of *baiA2* in the *bai* operon in *C. hylemonae. BaiA2* encodes a short chain dehydrogenase/reductase that is responsible for two steps in the 7α-dehydroxylation pathway. One of those steps is the conversion from cholyl-CoA to 3-oxo-cholyl-CoA in the oxidative arm, and the other is the conversion from 3-oxo-DCA to DCA in the reductive arm of the pathway (13, 50). However, it is important to note that while *C. hylemonae* lacks *baiA2* in the main *bai* operon*, baiA1* is present elsewhere in the genome under the control of a different promoter (12). *BaiA1* can also perform the conversion from cholyl-CoA to 3-oxo-cholyl-CoA (50, 51). When *baiA1* and *baiA2* from *C. scindens* VPI 12708 were expressed in *E. coli*, the substrate turnover (k_cat_) was higher for BaiA1 than for BaiA2 (50). It is also important to note that there may be other redundancies built in to the 7α-dehydroxylation pathway. When *baiN* was expressed in *E. coli*, it was capable of performing two conversions, that of 3-oxo-4,5-6,7-didehydro-DCA to 3-oxo-4,5-dehydro-DCA then to 3-oxo-DCA. More recently, the entire *bai* operon pathway was expressed in *C. sporogenes* and showed the same two steps being performed by different enzymes, BaiH and BaiCD respectively (13, 49). While both sets of enzymes are capable of performing these transformations, it is still unknown which one is preferentially utilized by *Clostridia* encoding the *bai* operon.

Given the complexity of the 7α-dehydroxylation pathway and the apparent redundancies built in, the production of DCA is likely very important for the commensal *Clostridia* encoding the *bai* operon. It was surprising to see relatively low MICs of all four commensal *Clostridia* against DCA (Figure 2), as this indicates that the organisms are producing something that is detrimental to them in a sufficient concentration. The reason these commensals produce DCA isn’t known, but dietary DCA has been shown to affect the microbiota in chickens and dietary CA supplementation in rats resulted in the outgrowth of *Clostridia* and an increase in DCA, suggesting the production of DCA may modulate the microbiome in a way favorable to these *Clostridia* (52, 53). While DCA, like other bile acids, has detergent-like properties, the specific mechanism of action of DCA or other bile acids involved in 7α-dehydroxylation, such as CDCA and LCA, against commensal *Clostridia* has yet to be elucidated. DCA and other bile acids have various effects on other bacteria found in the gut. In *Lactobacillus* and *Bifidobacteria*, DCA, as well as CA and CDCA, can inhibit growth by dissipating transmembrane electric potential and the transmembrane proton gradient (54). Bile acids, including DCA, can induce transcription of several genes responsible for DNA repair and recombination in *Escherichia coli, Salmonella typhimurium, Bacillus cereus* and *Listeria monocytogenes* (55). Genes responsible for maintaining the integrity of the cellular envelope were also upregulated in *B. cereus* and *L. monocytogenes*, indicating that bile acids such as DCA and CDCA damage the bacterial membrane and cellular DNA (55).

DCA and other bile acids such as CDCA and LCA have multiple effects on *C. difficile* as well. The presence of flagella, as well as the presence of the flagellar structural protein FliC, was significantly decreased when *C difficile* was challenged with DCA, CDCA and LCA, with LCA causing a near complete loss of flagellar filaments (24). CA did not have any significant effect on flagella in *C. difficile*, but cells challenged with CA, DCA or CDCA were significantly longer than the control cells, while LCA had no significant effect on cellular shape (24). While this indicates a potential mechanism for the inhibition of growth by DCA, CDCA and LCA, the mechanism for how these bile acids inhibit *C. difficile* toxin activity is still unknown (9, 24, 56).

*C. difficile* showed a very high tolerance to CA, with an MIC of 10 mM when grown in 100 µL of BHI (Figure 2). Interestingly, when *C. difficile* was grown in 200 µL of BHI in the growth kinetics assay, growth was observed at 10 mM CA, and complete inhibition only occurred at a concentration of 13 mM CA (Figure S3). CA also appeared to decrease the lag time of *C. difficile* when growth kinetics were analyzed (Figure S3), indicating that supplementation with CA could be beneficial to *C. difficile* growth as well as spore germination (22).

While all four commensal strains tested are capable of making DCA, only the *C. scindens* strains were capable of making enough to inhibit *C. difficile* (Figure 3A-B) in the *in vitro* assay performed. This is likely due to the higher amount of DCA produced when *C. scindens* was supplemented with 2.5 mM CA. Under the conditions tested, *C. hylemonae* did not produce DCA regardless of the amount of CA added to the media (Figure 3D) and no increased activity in the *bai* operon was observed when the media was supplemented with 2.5 mM CA or 7 mM CA. In another study, *C. hylemonae* did have increased expression of several genes in the *bai* operon, including *baiE* and *baiG* in a defined media supplemented with 0.1 mM CA. We tested expression of *baiG, baiE* and *baiN* in BHI media supplemented with 0.1 mM CA, and found no increase in expression (data not shown) (20). This indicates that regulation of the *bai* operon may be dependent upon changes in nutritional needs of the bacterium as well as the presence or absence of bile acids, and there may be strain specific differences in regulation. While *C. hiranonis* did produce DCA and had increased expression in *bai* operon genes (Figure 4C) under the conditions tested, supernatants from *C. hiranonis* were not able to inhibit *C. difficile* as it produced less DCA than either *C. scindens* strain (Figure 3C). This is likely due to the susceptibility of *C. hiranonis* to DCA, as it’s MIC for DCA was lower than that of *C. difficile* and the amount of DCA produced (0.78 mM) was the same as the MIC for DCA (Table S1). While we did not perform the inhibition assay with CDCA, which is converted to LCA by the *bai* operon, the expression of *baiG, baiE* and *baiN* was tested in *C. scindens* VPI 12708 and *C. hylemonae* cultures supplemented with 0.025, 0.25, and 1.25 mM of CDCA. No significant changes in expression were found in the presence of CDCA when compared to the no bile acid control (data not shown).

While the production of DCA is a factor in the inhibition of *C. difficile* by commensal *Clostridia* containing the *bai* operon, competition for nutrients may also play a role. Like most *Clostridia, C. scindens, C. hylemonae* and *C. hiranonis* encode enzymes that are used in Stickland fermentation, which is required for growth of *C. difficile* and several other *Clostridia* (20, 21, 36, 37). In particular, proline, which *C. difficile* is auxotrophic for, is one of the most efficient electron acceptors. Another amino acid that most commensal *Clostridia* can utilize for growth is hydroxyproline which can be converted to proline using the *hypD* gene, which is present in many *Clostridia* (57, 58). There is also the production of other inhibitory metabolites to consider, such as the tryptophan derived antibiotics 1-acetyl-b-carboline and turbomycin A that are produced by *C. scindens* ATCC 35704 and inhibit *C. difficile* (48). *C. difficile* is also capable of the production of inhibitory molecules. *C. difficle* ATCC 9689 produces proline-based cyclic dipeptides, which inhibit *C. scindens* ATCC 35704, *C. sordellii* and several other bacterial strains commonly found in the gut (48). As well as suggesting an important role for tryptophan, which is required for the growth of *C. difficile* and *C. scindens* ATCC 35704, this indicates a potential mechanism for the inhibition of commensal *Clostridia* when co-cultured with *C. difficile* (Figure 5) (21, 35).

Finally, there are some limitations to this study. As was seen with the expression of *baiE* and *baiG* in *C. hylemonae* (Figure 4), the type of media that is used for *in vitro* assays could affect expression of some genes. In addition, *in vitro* assays do not systematically mimic the *in vivo* environment, especially when studying a complex gut environment. In addition, due to the lack of genetic tools available for these commensal *Clostridia* at the time of the study, all assays were performed with wild type strains. This means that in the experiments assessing inhibition of commensal *Clostridia* by CA, the MIC values for *C. scindens* and *C. hiranonis* were likely affected by the conversion of some of the CA in the media to DCA in the assay. While 7α-dehydroxylation also transforms CDCA to LCA, only the inhibition of *C. difficile* by commensal *Clostridia* supplemented with CA was examined in this study due to solubility issues with LCA in media.

The genetic intractability of commensal *Clostridia* has made separating the inhibitory effects of 7α-dehydroxylation from other potentially inhibitory mechanisms such as nutrient competition or the production of antimicrobials difficult (48). However, a recently published CRISPR-Cas9-based method for constructing multiple marker-less deletions in commensal *Clostridia* shows great promise in assisting with this analysis (59). Future *in vitro* studies using defined media are also needed to further examine the role nutrient competition between *C. difficile* and commensal *Clostridia*. Additional *in vivo* studies are also needed to elucidate the direct mechanistic role 7α-dehydroxylation by commensal *Clostridia* plays in colonization resistance against *C. difficile.* Additional studies dissecting the regulation of the *bai* operon *in vitro* and *in vivo* and how this affects CDI will be important for future therapeutic interventions.

## Acknowledgements

We would like to thank Jason Ridlon for providing the commensal *Clostridia* strains. ADR was funded by the NCSU Molecular Biology Training Program T32 GM008776 through NIH. CMT is funded by the National Institute of General Medical Sciences of the National Institutes of Health under award number R35GM119438.

---

**Supplemental Figure 1: Optical density/Colony Forming Units conversion for commensal Clostridia and C. difficile.** Linear regression of OD_600_ plotted against CFU/mL for *C. scindens* **(A)**, *C. hiranonis* **(B)**, *C. hylemonae* **(C)** and *C. difficile* **(D)**.

**Supplemental Figure 2: Genome comparison of commensal Clostridia strains.** Whole-genome nucleotide BLAST alignment of *C. scindens* VPI 12708, *C. hiranonis*, and *C. hylemonae* against reference strain *C. scindens* ATCC 35704.

**Supplemental Figure 3: Cholate inhibits growth in Clostridia strains in a dose dependent manner.** Growth of *C. scindens* VPI 12708 **(A)**, *C. scindens* ATCC 35704 **(B),** *C. hiranonis* **(C)**, *C. hylemonae* **(D)**, and *C. difficile* **(E)** in the presence of multiple concentrations of CA along with a no bile acid control over 24 hr.

**Supplemental Figure 4: Deoxycholate inhibits growth in Clostridia strains in a dose dependent manner.** Growth of *C. scindens* VPI 12708 **(A)**, *C. scindens* ATCC 35704 **(B),** *C. hiranonis* **(C)**, *C. hylemonae* **(D)**, and *C. difficile* **(E)** in the presence of multiple concentrations of DCA along with a no bile acid control over 24 hr.

**Supplemental Figure 5: DCA inhibits C. difficile at a 2.5 mM concentration.** Inhibition of *C. difficile* after 24 hr of growth in media supplemented with 0.25 mM CA, 2.5 mM CA, 7.0 mM CA, 0.25 mM DCA or 2.5 mM of DCA. Experiments were run in duplicate and three biological replicates were performed. Inhibition was measured by comparison to a *C. difficile* control grown in media without bile acid supplementation. Statistical significance was determined by one-way ANOVA with Tukey used for multiple comparisons (*, p<0.05; **, p<0.01; ***, p<0.001; ****, p<0.0001).

**Supplemental Figure 6: C. difficile inhibits all commensal Clostridia strains except C. scindens ATCC 35704 in co-culture.** CFU/ml at 0 and 24 hr time points for monoculture and 1:1 competition with *C. difficile* for *C. scindens* VPI 12708 **(A)**, *C. scindens* ATCC 35704 **(B)**, and *C. hylemonae* **(C)**. Statistical significance was determined by one-way ANOVA with Tukey used for multiple comparisons (*, p<0.05; **, p<0.01; ***, p<0.001; ****, p<0.0001).

**Supplemental Table 1:** Log_2_ Fold Change for qRT-PCR with addition of CA

| Organism | Gene | 0.25 mM CA |  | 2.5 mM CA |  | 7.0 mM CA |  |
| --- | --- | --- | --- | --- | --- | --- | --- |
|  |  | Avg | SD | Avg | SD | Avg | SD |
| *C. scindens*<br>VPI 12708 | *baiG* | 9.75 | 0.15 | 10.89 | 0.21 |  |  |
|  | *baiE* | 10.32 | 0.72 | 11.94 | 0.53 |  |  |
|  | *baiN* | 0.10 | 0.29 | -0.15 | 0.15 |  |  |
| *C. scindens*<br>ATCC 35704 | *baiG* | 4.24 | 0.49 | 11.96 | 0.23 |  |  |
|  | *baiE* | 4.61 | 0.60 | 13.02 | 0.56 |  |  |
|  | *baiN* | 0.37 | 0.28 | 0.63 | 0.20 |  |  |
| *C. hiranonis* | *baiG* | 5.64 | 0.96 | 8.36 | 0.33 |  |  |
|  | *baiE* | 6.97 | 0.88 | 9.61 | 0.66 |  |  |
|  | *baiN* | 0.58 | 0.86 | 0.05 | 0.74 |  |  |
| *C. hylemonae* | *baiG* |  |  | -1.05 | 0.74 | -1.65 | 0.28 |
|  | *baiE* |  |  | -2.16 | 1.11 | -2.28 | 0.73 |
|  | *baiN* |  |  | -0.01 | 1.11 | 0.35 | 0.45 |

**Supplemental Table 2:** Minimum Inhibitory Concentration for CA and DCA (mM)

| Organism | Cholate (CA) |  | Deoxycholate (DCA) |  |
| --- | --- | --- | --- | --- |
|  | Average | SD | Average | SD |
| *C. scindens* VPI 12708 | 2.50 | 0.00 | 1.88 | 0.72 |
| *C. scindens* ATCC 35704 | 2.50 | 0.00 | 2.19 | 0.63 |
| *C. hiranonis* | 2.50 | 0.00 | 0.78 | 0.31 |
| *C. hylemonae* | 7.00 | 0.00 | 1.25 | 0.00 |
| *C. difficile* R20291 | 10.00 | 0.00 | 1.56 | 0.63 |

